# Disruption of lactate metabolism in the peripheral nervous system leads to motor-selective deficits

**DOI:** 10.1101/2022.06.29.497865

**Authors:** A. Joseph Bloom, Amber R. Hackett, Amy Strickland, Yurie Yamada, Joseph Ippolito, Robert E. Schmidt, Yo Sasaki, Aaron DiAntonio, Jeffrey Milbrandt

## Abstract

Schwann cells (SCs) myelinate and provide trophic support to axons in the peripheral nervous system (PNS) and disruption of SC cellular metabolism leads to demyelination and axon degeneration, both symptoms of peripheral neuropathies. The lactate shuttle hypothesis proposes that glycolytic support cells supply lactate to adjacent axons to sustain their high metabolic demands, a process that requires the interconversion of lactate and pyruvate via lactate dehydrogenase (LDH) in both SCs and neurons. To test this hypothesis in the PNS, we selectively knocked out the genes for both LDH enzymes, LDHA and LDHB, in motor neurons (MNs), sensory neurons (SNs), or SCs. Interestingly, motor axons and their synapses progressively degenerate when LDH is deleted from either MNs or SCs; however, defects in sensory axons or their terminals were not observed when LDH was excised from either SNs or SCs. Deletion of LDH in SCs also leads to a decrease in total ATP levels in peripheral nerves despite a marked accumulation of pyruvate and glycolytic intermediates, consistent with the failure of pyruvate to lactate conversion in SCs leading to energetic deficits in axons. These results support a model in which motor axons are more dependent on SC-derived lactate than are sensory axons, a specific dependency that suggests LDH and lactate shuttling influence the course of motor-dominated neuropathies such as ALS.

## Introduction

Schwann cells (SCs), the major support cells in the PNS, both myelinate and provide trophic support to axons^1^. Not surprisingly, disruption of metabolism specifically in SCs leads to defects in myelination and axon degeneration, both symptoms of peripheral neuropathy^2–8^. The lactate shuttle hypothesis proposes that glycolytic support cells actively produce lactate and transfer it to neurons to supplement their high metabolic demands^9^. Once transported from a SC into an axon, lactate is available for conversion to pyruvate and mitochondrial ATP production as well as macromolecular synthesis^10^. A few key pieces of evidence support this hypothesis. SCs store glycogen while axons do not^11^, and axonal and neuromuscular junction (NMJ) defects occur in mice that lack the lactate transporter MCT1 specifically in SCs^12,13^. However, these MCT1-deficient mice do not completely abrogate lactate shuttling from SCs to axons, perhaps due to other MCTs or other lactate transport mechanisms.

Lactate dehydrogenase (LDH) is the enzyme responsible for the interconversion of pyruvate and lactate. Functional LDH is a tetrameter comprised of combinations of two main subunits, LDHA and LDHB. A third isoform, LDHC, is expressed only in testis. While both LDHA and LDHB can catalyze the bidirectional interconversion of pyruvate and lactate, LDHA has a higher affinity for pyruvate than lactate, while LDHB has a higher affinity for lactate than pyruvate^14–16^. As the equilibrium of this interconversion is dependent on substrate concentrations, to abolish LDH activity it is necessary to delete both LDHA and LDHB^17^. Therefore, to determine the influence of SC lactate production on axonal maintenance, we deleted the *Ldha* and *Ldhb* genes specifically in motor neurons (MN), sensory neurons (SN), or SCs. In this manner, we prevented interconversion of pyruvate and lactate in either the axonal compartment or the surrounding glial cells, hence blocking potential lactate traffic between these compartments. We found progressive motor deficits in mice lacking LDH in either SCs or motor neurons. However, loss of LDH in sensory neurons or SCs did not cause significant sensory deficits. Motor deficits included functional abnormalities and pathological changes in both axons and NMJs. These results show that motor axons depend on lactate shuttling from SCs to meet their metabolic needs, while sensory axons appear to use other metabolic pathways to support their function. Based on this striking dichotomy of motor vs. sensory axon metabolic requirements, we hypothesize that LDH activity and lactate shuttling may contribute to motor-dominated neuropathies including ALS.

## Methods

### Animals

All animal experiments were performed under direction of institutional animal study guidelines at Washington University, St. Louis, MO. Floxed Ldha (*Ldha^F/F^*) mice were generated by breeding *Ldha*^tm1a(EUCOMM)Wtsi^ mice (EMMA, EM:05082, Jax Stock No: 030112)^18^ to mice that express FLP recombinase ACTB:FLPe B6;SJL (Jax Stock No: 003800)^19^. Floxed Ldhb (*Ldhb^F/F^*) mice were generated by breeding Ldhb^tm1a(KOMP)Wtsi^ mice (EMMA, EM:08936)^20^ to mice that express FLP recombinase. To generate Schwann cell, motor neuron or sensory neuron-specific LDH knockout mice, doubly mutant *Ldha^F/F^*; *Ldhb^F/F^* mice were bred to either MPZ-Cre^+ 21^, ChAT-Cre^+^ (Jax Stock No: 006410)^22^ or Na_v_1.8-Cre^+^ mice^23^ respectively. The Cre-expressing mouse strains were previously back-crossed to C57BL/6N >10 generations. All experimental mice were therefore of mixed C57BL/6N and B6Brd;B6Dnk;B6N-Tyr genetic background. Littermate controls consisted of Cre^-^ mice in all experiments. Except where specifically mentioned, male and female mice were used in approximately equal numbers in all experiments. Sciatic nerve crush experiments were performed as previously described^3,24^.

### Mouse Behavior

The inverted screen test was performed as previously described^24^. If a mouse did not fall within 120 s, then the time was recorded as 120□s. The tail flick assay was performed as previously described ^24,25^. The Von Frey assay was performed as previously described^24,26^. To begin, mice were stimulated with filaments (Semmes-Weinstein monofilaments, Stoelting) for 2 s in the center of the foot between the footpads starting at 0.32□g of force. Subsequently, the up-down method^27^ was used. Mice were stimulated four times after the first change in withdrawal for each foot separately. The withdrawal threshold was calculated and the values for each foot were averaged.

### Nerve electrophysiology

The compound muscle action potential (CMAP) and sensory nerve action potential (SNAP) were acquired as previously described^26,28^ using a Viking Quest electromyography device (Nicolet). Mice were anesthetized using isoflurane, then for CMAP, the recording electrodes were placed in the footpad and stimulating electrodes in either the ankle or sciatic notch. For SNAP recording, electrodes were inserted subcutaneously into the tail. The recording electrode was inserted at the base of the tail, the stimulating electrode was inserted 30 mm distal to the recording electrode, and the ground was inserted between the stimulating and recording electrodes. Supramaximal stimulation was used for CMAP and SNAP.

### QPCR

QPCR was performed as previously described^29^. Primer sequences were as follows: β-Actin (Forward GACATGGAGAAGATCTGGCA; Reverse GGTCTCAAACATGATCTGGGT), LDHA (Forward TGTCTCCAGCAAAGACTACTGT; Reverse GACTGTACTTGACAATGTTGGGA), and LDHB (Forward: CATTGCGTCCGTTGCAGATG; Reverse GGAGGAACAAGCTCCCGTG).

### LDH flux assay

LDH flux assay was performed as previously described^30^. Sciatic nerves were dissected from 3-month-old isofluorane anesthetized mice, epineurium removed in PBS, nerves cut into 4 approximately equal segments, and the nerve segments placed in uncoated 12 well plates with Neurobasal Media without glucose or pyruvate (ThermoFisher Scientific A2477501) supplemented with 10 mM ^13^C_6_-glucose for 24 hours. Samples were removed, washed with PBS, snap frozen on dry ice and stored at −80°C. Tissue was sonicated in 50% MeOH and the extract was evaporated under nitrogen gas and were derivatized with 14 μL of 20 mg/mL methoxyamine HCl in pyridine and incubated at 38°C for 90 min. Immediately following the 90 min incubation, 56 μL of MTBSTFA with 1% t-BDMCS were added and samples were incubated at 38°C for 30 min^30,31^. Derivatized samples were run on an Agilent 7890A GC coupled to an Agilent 5975C MS and data was acquired and analyzed in MSD ChemStation E.02.02.1431 (Agilent). The GC temperature program was set to: initial temperature: 60°C, hold for 0.65 min; temperature rate 1: 11.5°C/min to 201°C, hold for 0.65 min; temperature rate 2: 3.5°C/minute to 242°C, hold for 0 min; temperature rate 3: 16°C/min to 300°C, hold for 7 minutes. Data on all isotopologues was collected in SIM mode. Isotopologue peak areas in each sample were normalized in FluxFix^32^ to isotopologues of unlabeled samples to adjust for the natural abundance of each isotopologue.

### Steady state metabolomics

Steady state metabolite measurement was performed as previously described ^33^, for 3 and 6-month-old WT and SCKO mice. Briefly, sciatic nerves from 3-month-old mice were quickly dissected, the epineurium was removed in ice cold PBS, snap frozen on dry ice, and stored at −80°C. Metabolites were extracted by homogenizing frozen sciatic nerves in 50% MeOH (100 μL per nerve) using sonication and incubated 10 min on ice. The homogenates were centrifuged (10,000 × *g* for 15 min), and the supernatants containing small metabolites and the precipitates containing proteins were collected. The metabolite-containing fractions were further purified with chloroform (50 μL) extraction. The precipitates containing proteins were solubilized by adding 0.1% SDS solution (100 μL per nerve), and the protein content was determined using BCA protein assay kit (Thermo Fisher Scientific). The aqueous phase (90 μl) was lyophilized and stored at −20°C until analysis. Lyophilized samples were reconstituted with 5 mM ammonium formate.

For the measurement of metabolites including NAD^+^ and ATP, samples were injected into a C18 reverse phase column (Atlantis T3, 2.1 × 150 mm, 3 μm; Waters) using HPLC (Agilent 1290 LC) at a flow rate of 0.15□ml/min with 5 mM ammonium formate for mobile phase A and 100% methanol for mobile phase B. Metabolites were eluted with gradients of 0–10□min, 0–70% B; 10–15□min, 70% B; 16–20□min, 0% B. The metabolites were detected with a triple quad mass spectrometer (Agilent 6470 MassHunter; Agilent) under positive ESI multiple reaction monitoring (MRM) using parameters specific for each compound (NAD^+^, 664>428, fragmentation (F) = 160 V, collision (C) = 22 V, and cell acceleration (CA) = 7 V: ATP 508>136, F=90 V, C=30 V, CA = 27 V). Serial dilutions of standards for each metabolite in 5□mM ammonium formate were used for calibration. Metabolites were quantified by MassHunter quantitative analysis tool (Agilent) with standard curves and normalized by the protein. Alternative metabolite analysis targeting metabolites in the glycolysis pathway was performed as instructed (Metabolomics dMRM Database and Method, Agilent Technologies). Samples were chromatographically separated using a reverse phase column (Zorbax RRHD Extend-C18, 2.1 × 150 mm, 1.8 μm, Agilent Technologies) with mobile buffer A (3% MeOH, 10 mM tributylamine, 15 mM acetic acid) and 0%–99% gradient of buffer B (100% MeOH, 10 mM tributylamine, 15 mM acetic acid). Metabolites were analyzed by mass spectrometer (6470, Agilent Technologies) using dynamic multiple reaction monitoring. The compound signatures, including retention time and qualifying ions, are predetermined in the dynamic multiple reaction monitoring database and data analysis tool automatically detect these compounds. The relative amount of each metabolites was calculated and normalized by the amount of protein.”

### Nerve structural analysis using light and electron microscopy

Sciatic, sural, and the motor branch of the femoral nerve were processed as previously described with modifications^24,26,29^. First, nerves were pinned onto a Sylgard plate to maintain orientation while incubated overnight at 4°C in 3% glutaraldehyde in 0.1 m□PB (Polysciences). Next, nerves were sutured to demarcate the most distal region. They were washed with PB and incubated overnight at 4°C in 1% osmium tetroxide (Sigma-Aldrich). After washing, nerves were dehydrated in a serial gradient of ethanol from 50 to 100%, incubated in 50% propylene oxide/50% ethanol then 100% propylene oxide. Nerves were then placed in Araldite resin solution/propylene oxide solutions overnight in the following ratios: 50:50, 70:30, 90:10. Each nerve was cut in half and a block was made for the most distal and proximal portions. These fragments were embedded in 100% Araldite resin solution (Araldite: DDSA: DMP30; 12:9:1; Electron Microscopy Sciences) and baked overnight at 60°C. The blocks were cut using Leica EM UC7 Ultramicrotome to generate semithin sections 400–600□nm thick which were dried onto microscopy slides. For electron microscopy, blocks were cut into 300–400□nm sections and collected on copper grids. The grids were stained with uranyl acetate and lead citrate before imaging using a transmission electron microscope (JEOL1200).

### Toluidine blue staining and quantification

Prewarmed slides with 400–600□nm sections were stained for 1 minute with 1% toluidine blue solution (1% toluidine blue, 2% borax), vigorously washed with water, acetone, then xylene. Slides were mounted with Cytoseal XYL (Thermo Fisher Scientific). Whole nerves were imaged (sciatic at 10×, femoral at 20×, and sural at 40× magnification) and 3 random fields were obtained using a 100× oil-immersion lens. All of the myelinated axons in each image were counted and categorized into one of six categories—*normal, degenerated, hyper-myelinated, naked, onion bulb*, or *regenerated cluster*—by a trained observer blinded to genotype. Counts were normalized per square millimeter and averaged over three fields. Cell nuclei were similarly counted. To determine axon diameter and G-ratio, 40 axons per mouse were measured and analyzed using ImageJ. Degenerated axons were excluded from axonal diameter and G-ratio calculations.

### Muscle Fiber Cross-sectional area

Tibialis anterior muscles were dissected from mice transcardially perfused with 4% paraformaldehyde in PBS (PFA), postfixed for 1 hour at room temperature, then incubated for 48□hr in 30% sucrose in PBS. Samples were embedded in O.C.T. and sectioned at 10□μm using a cryostat. Sections were mounted on positively charged slides and stored at −20°C. Slides were postfixed in ice-cold acetone for 5 minutes, washed with PBS, blocked with 5% normal goat serum in PBST, then incubated in primary antibody (Laminin, Sigma (#L9393))

### NMJ staining and analysis

Lumbrical muscles dissected from mice transcardially perfused with 4% PFA were post-fixed in 4% PFA overnight and washed thrice for 15 min in PBS. Samples were permeabilized with 2% TritonX-100/PBS (PBST) for 30 min, and blocked with 4% bovine serum albumin (BSA) in 0.3% PBST for 30 mins at room temperature. Muscles were incubated with anti-SV2 (Developmental Studies Hybridoma Bank AB2315387, 1:200) and anti-2H3 (Developmental Studies Hybridoma Bank AB2314897, 1:100) for 48 hr at 4°C. After incubation with primary antibodies, muscles were incubated with FITC rabbit anti-mouse IgG (Invitrogen A21121, 1/400) overnight at 4°C. The samples were then incubated with CF 568 conjugated α-bungarotoxin (BTX) (Biotium 00006, 1:500) for 2 hr at room temperature. To analyze NMJ morphology, z-stack images using a confocal microscope (Leica DFC7000T) were obtained. Maximal intensity projection images were reconstructed, and post-synapse volume and terminal axon diameter were analyzed using Imaris software.

### Motor neuron quantification

Paraffin embedded spinal cords were sectioned at 6 microns. After deparaffinization, antigen retrieval in citrate buffer, pH 6.0 using a pressure cooker for 3 min. Tissues were incubated with rabbit CHAT antibody 1:300 (Millipore Cat #AB143) overnight at 4C, followed by goat anti-rabbit Cy3 1:500 1hour at RT.

CHAT positive cells were counted from 10x images in the ventral horn of the lumbar region of the spinal cord from 3-6 sections per animal.

### Intraepidermal nerve fiber quantification

For intraepidermal nerve fiber (IENF) staining and quantification, footpad skin was dissected and prepared as previously described^26^. The footpads were imaged on a Leica DMI 4000B confocal microscope using a 20X oil objective and a Leica DFC 7000-T camera. Z-stacks were acquired through the whole sample, max projected, then exported for quantification. IENF density was quantified as the number of PGP9.5^+^ axons that crossed the basement membrane normalized to the length of the basement membrane. IENF densities were averaged in three separate sections for each animal. Imaging and analysis were performed blinded to genotype.

### Statistical Analysis

Sample number (n) is defined throughout as the number of animals that were independently manipulated and measured. Histology, enzyme assays, and qPCR assays were always performed in 3 or 4 mice, whereas behavior and electrophysiology tests were performed with >5 animals per cohort. Statistics were performed using either Prism (GraphPad) or R.

## Results

### Motor neurons but not sensory neurons require LDH to maintain axonal integrity

To be utilized as an energy source to generate ATP, lactate must first be converted to pyruvate by LDH. Therefore, to probe the lactate shuttle hypothesis in the PNS, we generated peripheral neuron specific LDH knockout mice using the Cre-lox system, mating floxed *Ldha/Ldhb* animals to two different mouse lines: 1) ChAT-Cre^+^ mice, which express Cre in cholinergic neurons including α-motor neurons ^22^, or 2) Na_v_1.8-Cre^+^ mice which express Cre in ~75% of their dorsal root ganglion (DRG) neurons, including most nociceptors and ~40% of A-fibers ^23^. Mice lacking both LDHA and LDHB in motor neurons (MNKO mice) using ChAT-Cre or in sensory neurons (SNKO mice) using Na_v_1.8-Cre produced mice that were viable, fertile, and normal in appearance. To determine whether LDH activity is necessary for neuron function and axon maintenance, we subjected MNKO and SNKO mice to batteries of behavioral and electrophysiological tests to assess their respective motor and sensory function over time. Notably, MNKO mice showed significant weakness at six months of age that progressed to more severe motor dysfunction at twelve months (**Fig. 1A**). MNKO did not display a significantly reduced compound muscle action potential (CMAP) amplitude or nerve conduction velocity (NCV) at any age (**Fig. S1**).

**Figure 1:**
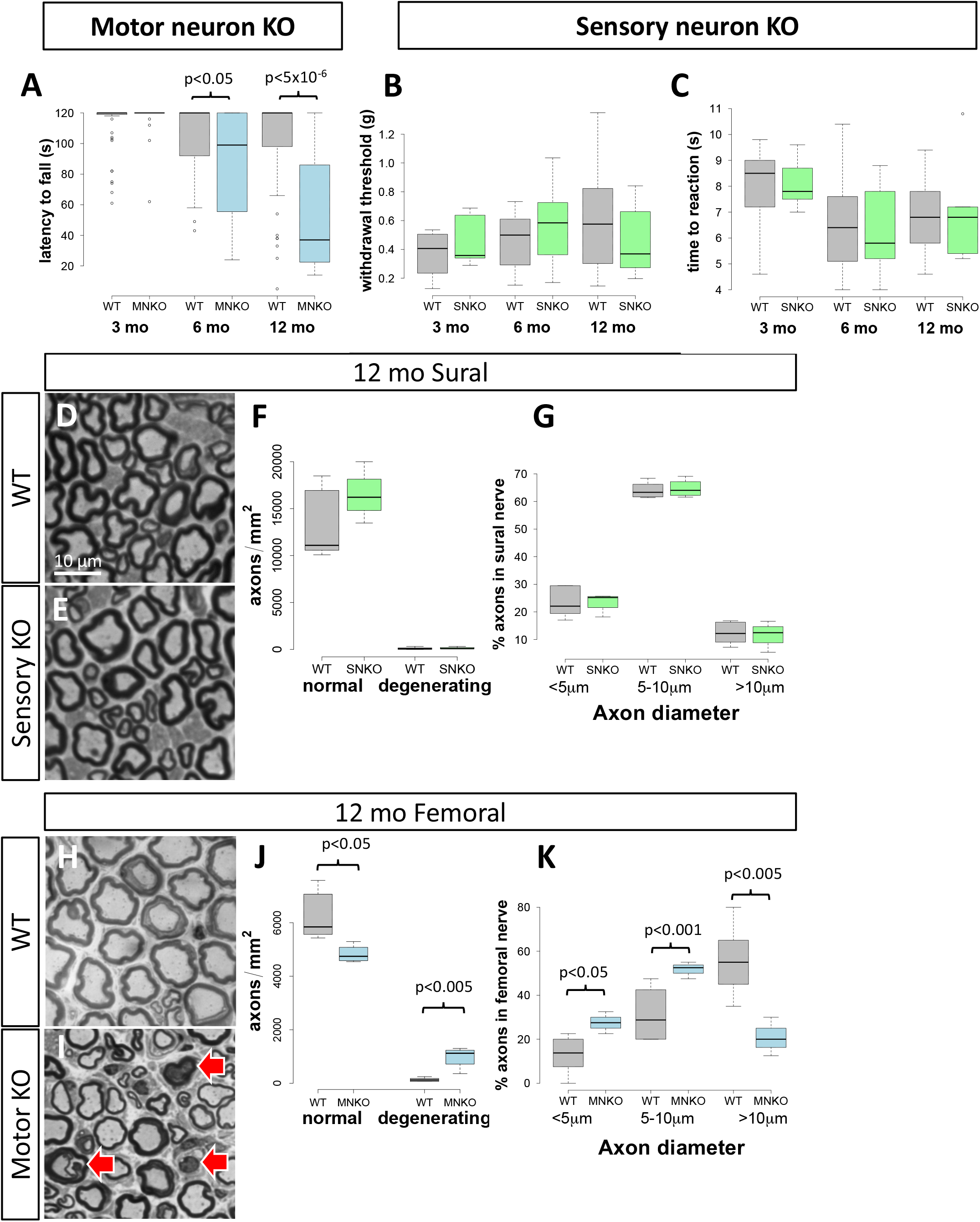
Deletion of LDH in motor but not sensory neurons leads to neuropathy. Motor function in motor neuron-specific LDH KO (MNKO) and wildtype (WT) control mice was assessed by measuring latency to fall from an inverted screen **(A)**. Mechanical sensation was assessed in sensory neuron-specific LDH KOs (SNKO) and controls by the Von Frey test **(B)** and heat sensation was assessed by the tail flick withdrawal assay **(C)**. Representative toluidine blue-stained sural (sensory) nerves from 12 month-old control and SNKO mice **(D**,**E)** and femoral (motor) nerves from MNKO mice **(H**,**I)** with controls. Red arrows indicate aberrant and degenerated axons. Graphs show the density of normal axons and degenerating axons, and the distribution of axon diameters in SNKO and control sural nerves **(F,G)**, and MNKO and control femoral nerves **(J,K)**.

In striking contrast, no differences in response to thermal or mechanical stimuli were observed in SNKO mice (**Fig. 1B,C**). The sensory nerve action potential (SNAP) amplitude and conduction velocity in the tail (**Fig. S1**) were not significantly different between genotypes, indicating there is no impairment in myelination or reduction in the number of myelinated axons in the sensory nerves of SNKO mice.

We next analyzed the morphology of the predominantly sensory sural nerve^34^ and the motor branch of the femoral nerve in 12 month old LDH-deficient mice. SNKO mice showed no signs of axonal degeneration or obvious defects in nerve morphology **(Fig. 1D-G)**, consistent with the lack of sensory functional deficits. In contrast, MNKO mice had fewer total axons in the femoral nerve, with particularly large losses of larger diameter axons, and an increased number of degenerated axons **(Fig. 1H-K)**. In the PNS, axon degeneration stimulates compensatory axon regeneration. While regenerating axon clusters and onion bulbs were not observed in femoral nerves of control animals (n=6), both were found frequently in MNKO femoral nerves (n=4, 454±81 regenerating clusters/mm^2^, p<1×10^−4^; 558±180 onion bulbs/mm^2^, p<5×10^−9^), findings consistent with ongoing axon degeneration. Consistent with motor axon loss and motor function deficits, we observed significant neuromuscular junction denervation of lumbrical muscles in the hind paws in 12-month-old MNKO mice with many unoccupied endplates **(Fig. 2A-C)**. Finally, to determine whether the observed axon degeneration was accompanied by neuron death, we counted ChAT^+^ motor neurons in the spinal cord of MNKO mice and found no significant difference from wildtype or SCKO mice at 12 months of age **(Fig. 2D)**. In sum, these results strongly suggest that motor but not sensory neurons are dependent on cell-autonomous LDH activity to maintain their axons.

**Figure 2:**
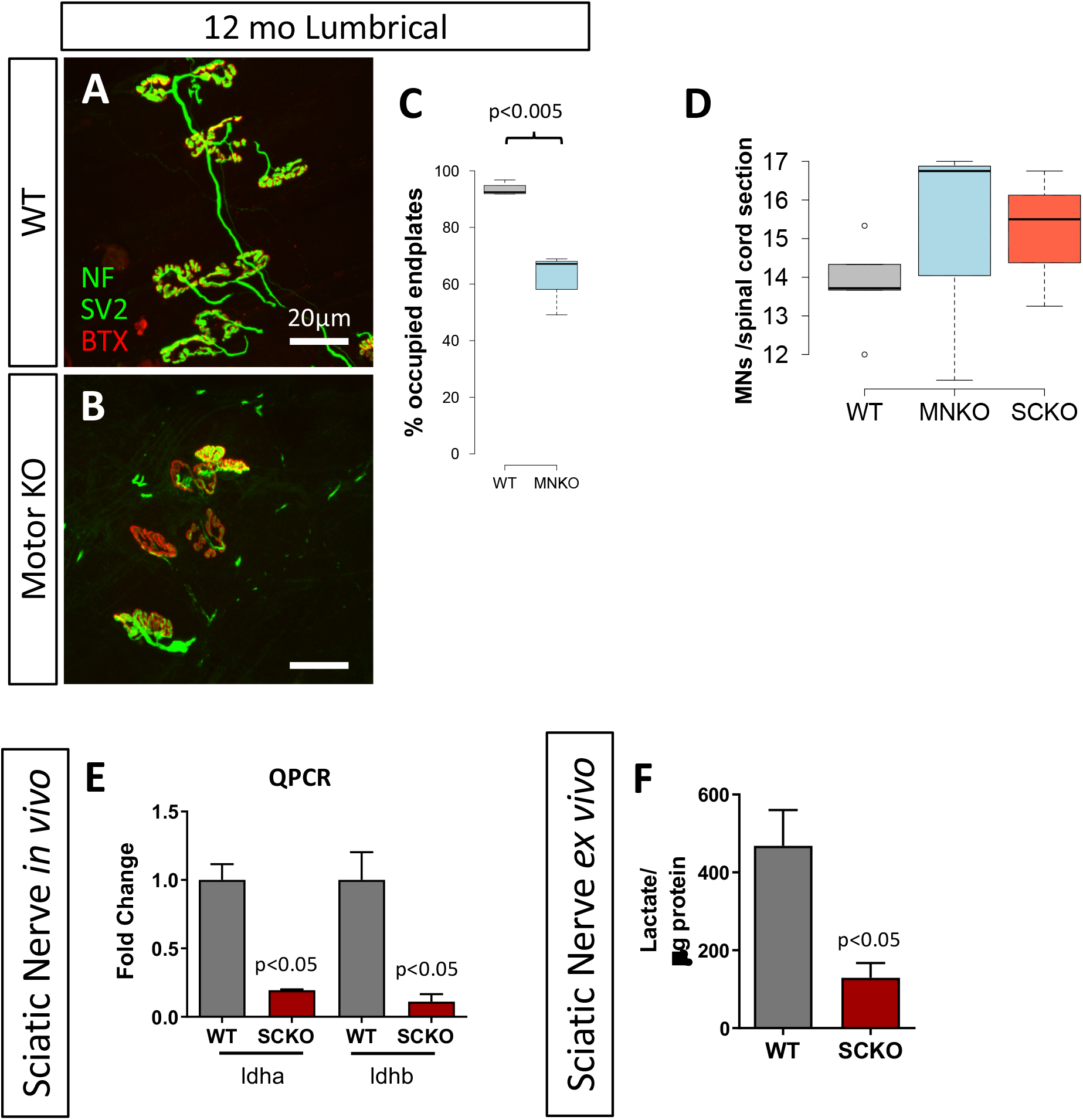
Deletion of LDH in motor neurons causes muscle denervation, LDH deletion from motor neurons or Schwann cells does not cause motor neuron loss in the spinal cord. NMJs on 12-month-old control and MNKO lumbrical muscles stained in green to detect the synaptic vesicle marker synaptic vesicle glycoprotein 2A (SV2) and axon marker neurofilament medium chain (NF) and in red with the post-synaptic endplate ligand bungarotoxin (BTX) **(A,B)**. Percent of the lumbrical muscle endplates innervated by apposed pre-synaptic structures **(C)**. Average ChAT+ motor neurons per section of spinal cords counted in 12-month old MNKO, Schwann cell KO (SCKO) and control mice **(D)**. Sciatic nerves from 3-month-old SCKO mice show reduced *ldha* and *ldhb* gene expression by QPCR **(E)**. Total lactate detected in 3-month-old sciatic nerves cultured 24 hours *ex vivo* **(F)**.

### LDH deficiency in Schwann cells leads to reduced pyruvate to lactate conversion and decreased ATP production in peripheral nerves

According to the lactate shuttle hypothesis, glial cells must produce lactate, which requires the activity of LDH, and then shuttle it to neurons/axons to support their function. To test this hypothesis in the PNS, we generated Schwann cell-specific LDHA/B KO mice by mating *Ldha/Ldhb* floxed mice to MPZ-Cre mice^21^. These mice express Cre in SC precursor cells beginning at E14, well before the initiation of myelination, which occurs postnatally. These LDH-deficient SC (SCKO) mice are viable, fertile, and appear overtly normal. The deletion of *Ldha* and *Ldhb* in SCs was detected by qPCR analysis **(Fig. 2E)**. Sciatic nerve segments from SCKO mice were cultured *ex vivo* and total lactate levels were dramatically reduced **(Fig. 2F)**, indicating that much of sciatic nerve lactate production occurs in SCs, as opposed to other cellular components of the nerve.

To assess effects on nerve lactate production resulting from loss of LDH in Schwann cells, we performed a metabolic flux assay using an *ex vivo* sciatic nerve culture system (**Fig. S2**). Nerves from wildtype and SCKO mice were grown in media supplemented with ^13^C_6_-labeled glucose for 24 hr and metabolites were extracted. All possible lactate isotopologues derived from ^13^C_6_-labeled glucose were measured by GC-MS. The ^13^C labeling pattern in lactate indicated that >70% of lactate in wildtype nerve is derived via its direct synthesis from glucose via glycolysis. As expected, the loss of LDH in SCs overwhelmingly blocks this pathway (**Fig. S2**), such that only small amounts of ^13^C_6_-labeled lactate are produced in SCKO nerve.

To further explore how loss of LDH in SCs alters nerve glycolysis, we measured the abundance of glycolytic metabolites in wildtype and SCKO sciatic nerves using LC-MS. We found that glyceraldehyde 3-phosphate, dihydroxyacetone phosphate, and phosphoenol pyruvate are all significantly increased in the SCKO model **(Fig. 3B-D)**. Pyruvate levels are robustly increased, with a 4-fold difference between SCKO and wildtype nerves at 3 months, and a 6.7-fold difference at 6 months of age **(Fig. 3E)**. We also observed a trend toward increased levels of pentose phosphate metabolites such as ribose 5-phosphate and xylose 5-phosphate, and purine metabolism products such as inosine, hypoxanthine, xanthine, and adenosine in SCKO nerves **(Fig. S3)**. Finally, we observed significantly reduced levels of ATP in SCKO nerves **(Fig. 3F)**. Overall, these data indicate that LDH actively converts pyruvate into lactate in SCs and that deleting LDH in SCs leads to a loss of lactate and an accumulation of pyruvate, glycolytic metabolites, and other metabolic intermediates (**Fig. 3A**).

**Figure 3:**
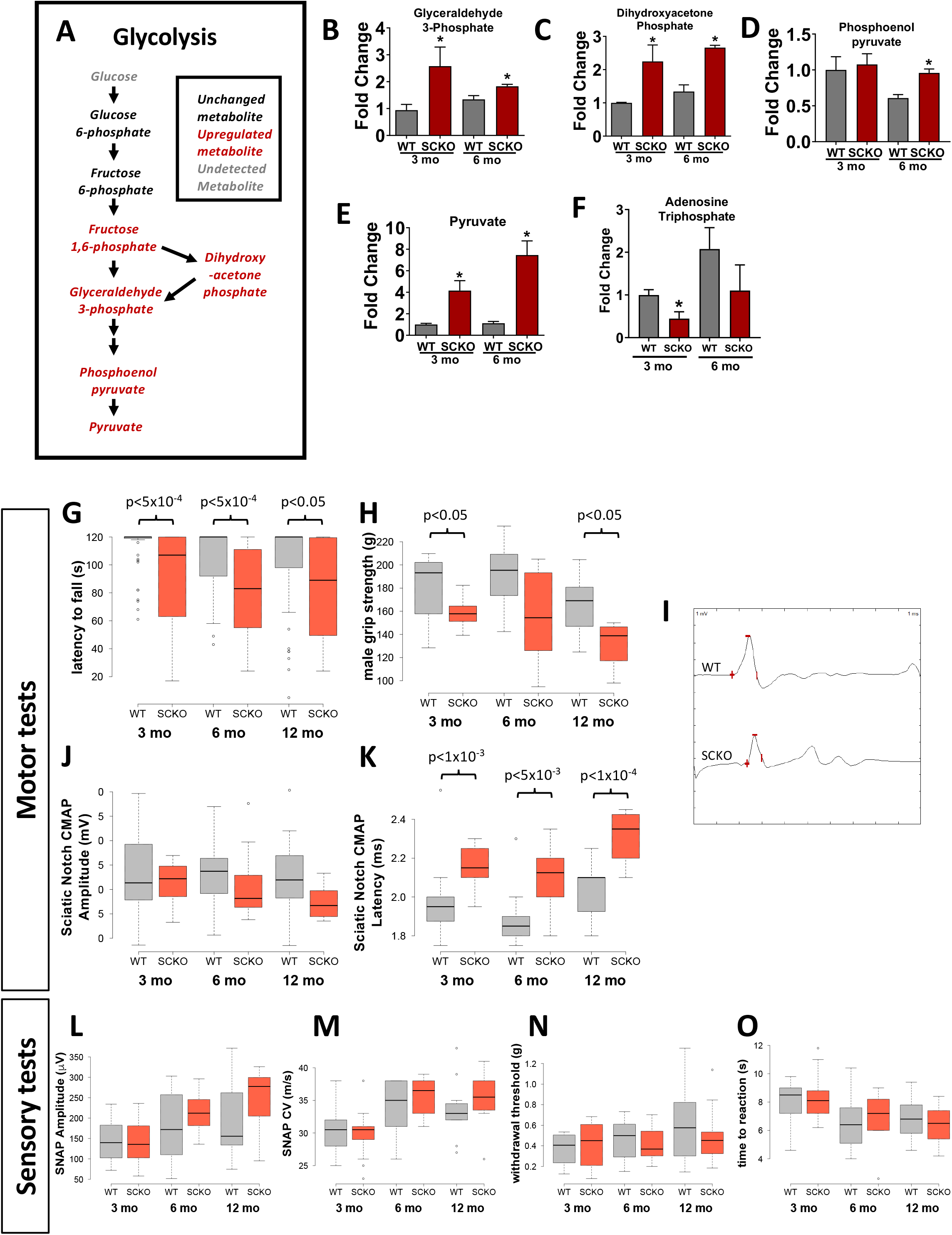
Deletion of LDH in Schwann cells alters glycolytic metabolites in sciatic nerves and causes motor but not sensory defects. Metabolic analysis of glycolysis in 3-month-old SCKO sciatic nerves **(A-F).** Metabolites in gray were undetected, metabolites in black were unchanged, and metabolites in red were upregulated. ATP **(F)** was significantly reduced. n=3-5 mice, *p<0.05 using a Student’s t-test between genotypes at each time point. Behavioral and electrophysiological tests were performed to assay motor **(G-K)** and sensory **(L-O)** function at 3, 6, and 12-month old mice, including inverted screen **(G)** forelimb grip strength in male mice **(H),** measurement of Compound Muscle Action Potential (CMAP) at the Sciatic Notch Amplitude **(I,J)** and latency **(K)**, Sensory Nerve Action Potential (SNAP) amplitude and conduction velocity (CV) measured from the tail **(L,M)**, mechanical sensitivity assessed via Von Frey test **(N)** and thermal sensitivity determined by latency to respond to submerging the mouse’s tail in a 50°C water bath **(O)**, n≥6 mice.

### Deletion of LDH in Schwann cells leads to progressive motor but not significant sensory neuropathy

To examine the functional consequences of SC LDH loss, we performed motor and sensory behavioral tests. We found that SCKO mice display a progressive diminution of strength similar to that observed in the MNKO model (**Fig. 3G,H)**, strongly supporting the findings presented above that motor axons are dependent on lactate to maintain proper function. The sciatic nerve CMAP amplitudes were also reduced in 12 month old SCKO mice (**Fig. 3I,J**), suggesting motor axon loss, while CMAP latency was increased (**Fig. 3K**). NCV was unchanged in SCKO mice at 12 month of age (**Fig. S1**). By contrast, both behavioral and electrophysiological measures of sensory function appear normal in 12-month-old SCKO mice (**Fig. 3L-O**).

We next examined the structure of peripheral nerves in SCKO mice to determine the basis of the motor deficits. In young animals (P7 and P21), we found no differences in the number of myelinated axons, myelin thickness, axonal diameter distribution, or number of SC nuclei between SCKO and control mice **(Fig. S4)**, suggesting that PNS development proceeds normally despite lack of LDH activity in SCs. Surprisingly, we also observed no difference in the speed or extent of remyelination after nerve injury in 2-month old SCKO mice (**Fig. S5**). In SCKO mice at 3 months of age, the distal sciatic nerve—which contains both motor and sensory axons—remains grossly normal, although tomacula and degenerating axons are present **(Fig. 4 & 5)** as are hypermyelinated axons [SCKO: 408±129/mm^2^ (n=5) vs. WT: 50±37/mm^2^ (n=4), p<0.05]. In older SCKO mice, there is evidence of segmental demyelination in some sectors of the distal nerve. By 12 months of age most axons in these sectors display thinner myelin (i.e. higher g-ratio, **Fig. 5A**) and we observed frequent morphological hallmarks of peripheral neuropathy (n≥3) including tomacula, degenerating axons (**Fig. 5C**), naked axons (105±12/mm^2^, p<0.005), supernumerary SCs resembling onion bulbs (908±115/mm^2^, p<0.005), and regenerating clusters (several axons wrapped by a supernumerary SC, 344±55/mm^2^, p<0.01) **(Fig. 4 & 5)**. Degeneration is substantially less severe in the proximal portion of the nerve (**Fig. S6)**, suggesting that transport of metabolites from the cell body bolsters axonal health when SC support is withdrawn.

**Figure 4:**
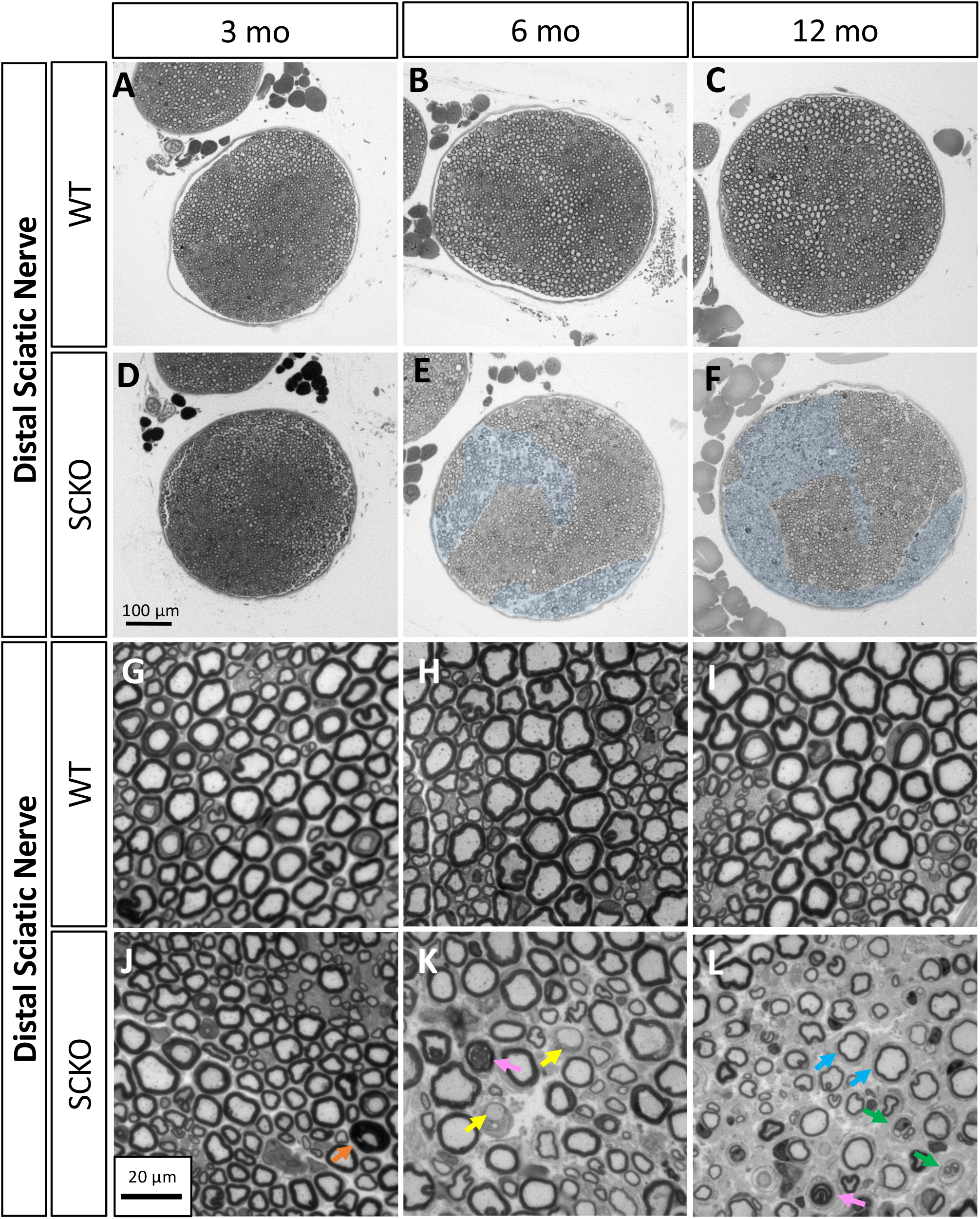
Sciatic nerves from SCKO mice have defects in a subpopulation of myelinated axons. Distal sciatic nerve cross-sections stained with Toluidine blue at 3, 6, and 12 months of age in WT and SCKO mice. Blue pseudo-coloring indicates affected sectors **(A-L)**. At 3 months of age, SCKO nerves appear grossly normal, except for the presence of tomacula **(J** orange arrow). At 6 months, affected sectors in SCKO nerves have naked axons **(K** yellow arrows) and degenerated axons **(K** pink arrow). At 12 months, most axons in affected sectors of SCKO nerves have defects, including onion bulbs **(L** blue arrows), regenerating clusters **(L** green arrows), and degenerating axons **(L** pink arrow).

**Figure 5:**
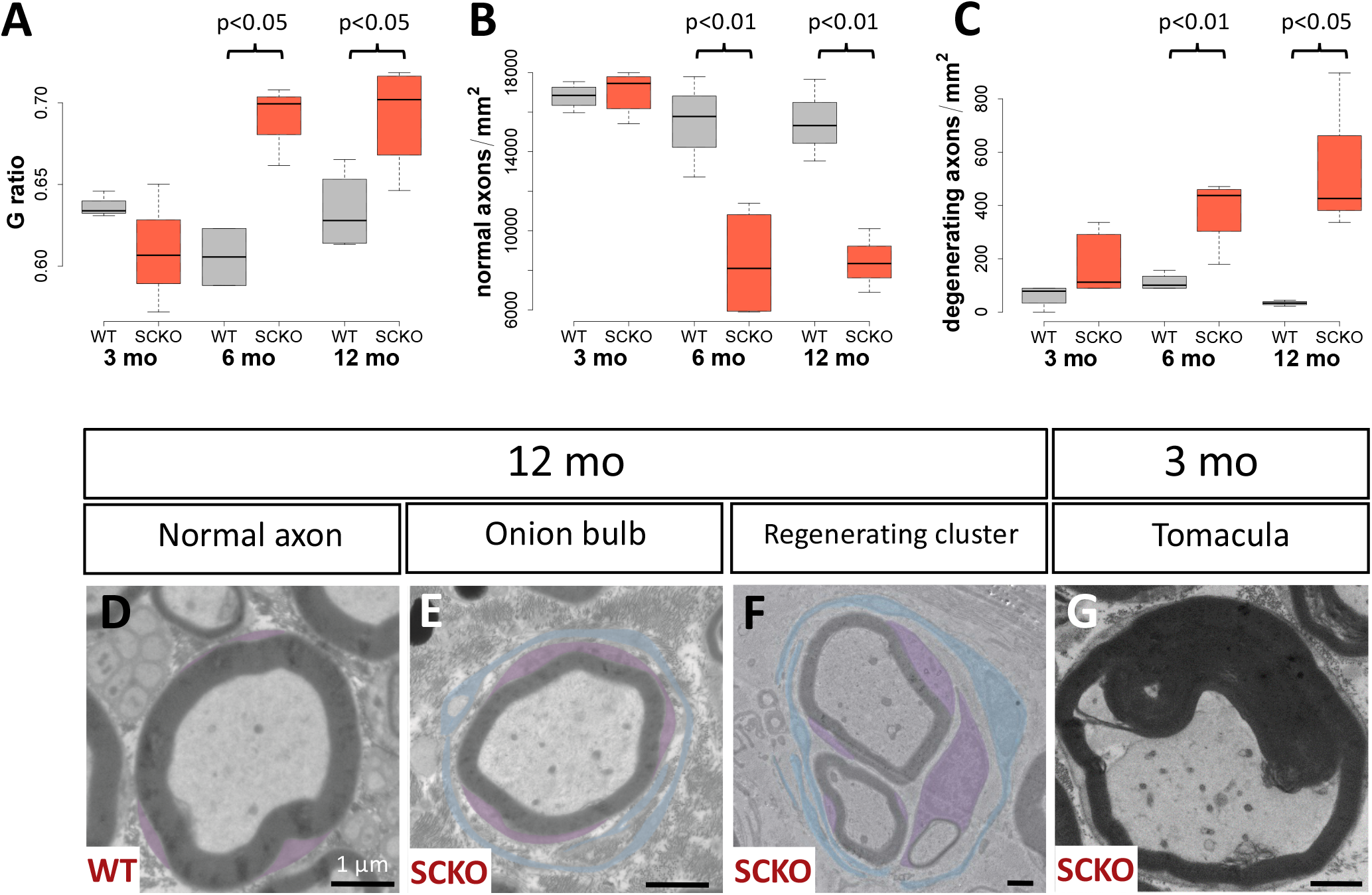
Detailed histomorphometry reveals progressive axon degeneration in SCKO mice. Average G-Ratio **(A)**, density of normal axons **(B)**, and degenerating axons **(C)** in sciatic nerves of WT and SCKO mice at 3, 6 and 12 months of age, n = 3-5 mice. Representative images of a normal axon **(D)**, an onion bulb **(E)**, a regenerating cluster **(F)**, and a tomacula **(G)**.

Tellingly, sectors of the SCKO sciatic nerve that demonstrate evidence of axonal degeneration mostly contain large diameter axons consistent with distribution of motor fascicles in this nerve^35^. Therefore, to probe the apparent motor-specificity of these defects, we again focused on the primarily sensory sural nerve and the motor branch of the femoral nerve in 12 month old mice. We observed no apparent abnormalities in the sural nerve **(Fig. 6A-D)** but many morphological changes in the femoral nerve (**Fig. 6E-H**). No differences in the density of axons, number of degenerating axons (**Fig. 6I**) or axonal diameter (**Fig. 6J, K**) are detected in the sural nerves of SCKOs, whereas their femoral nerves contain significantly fewer axons (**Fig. 6L**) and those axons that do remain are smaller in diameter (**Fig. 6M-N**). Femoral nerves from 12-month old SCKO animals contain significantly more degenerating axons **(Fig. 6L)**, regenerating clusters (737±124/mm^2^ (n=3) vs. 0 in controls (n=6), p<5×10^−4^)), onion bulbs (1,174±43/mm^2^ vs. 0, p<5×10^−9^), and naked axons (67±11/mm^2^ vs. 0, p<1×10^−4^). Regenerating clusters and onion bulbs are hallmarks of peripheral neuropathy and occur in CMT1 (demyelinating), CMT2 (axonal), and CMTX (X-linked) ^36–38^. Overall, these results are consistent with repetitive axon demyelination and remyelination in the femoral and sciatic nerves of SCKO mice.

**Figure 6:**
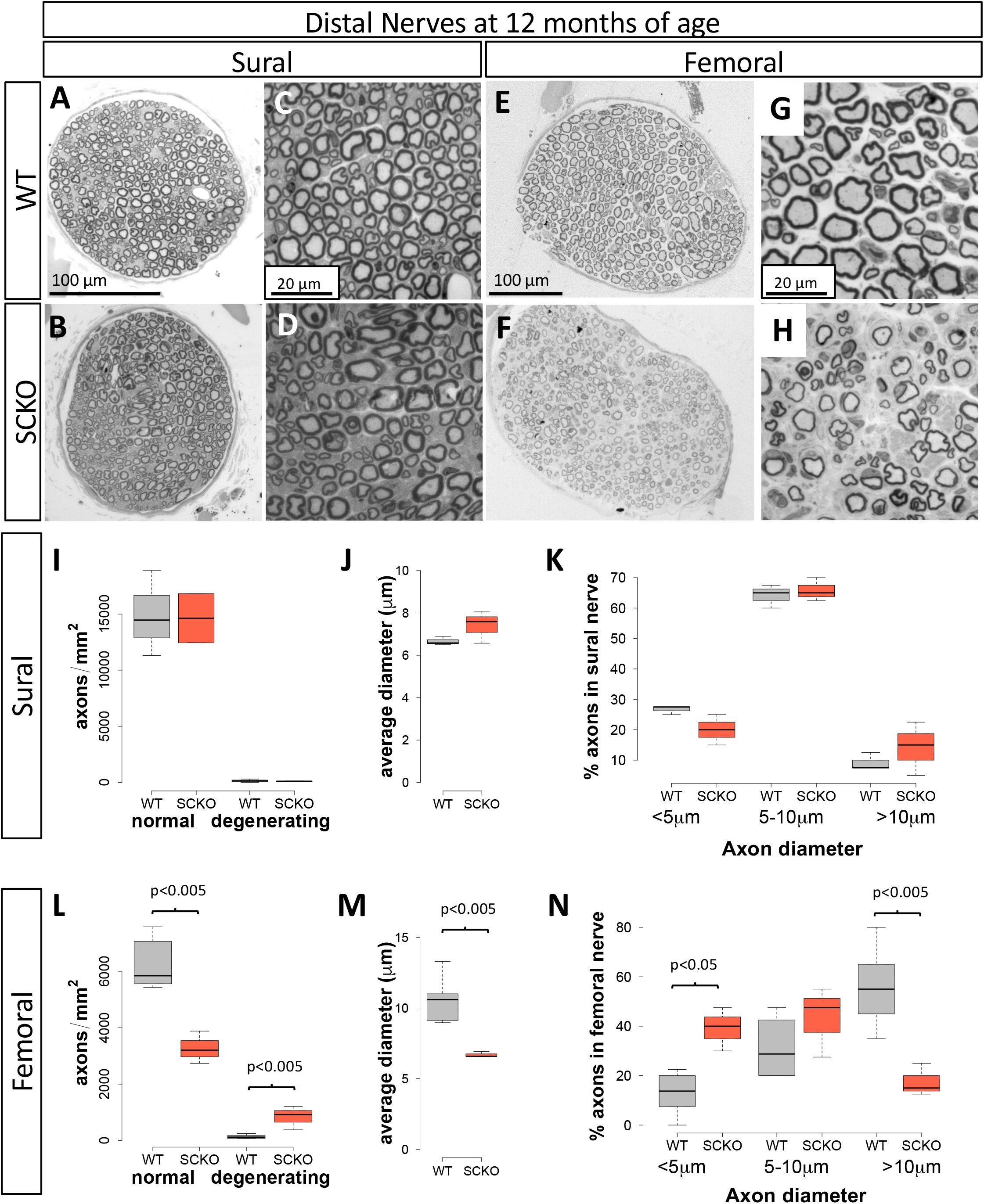
Motor nerves show more severe defects than sensory nerves in SCKO mice. Toluidine blue-stained (sensory) sural and (motor) femoral nerves from 12-month-old WT and SCKO mice **(A-H)**. Blue arrows indicate onion bulbs, green arrows indicate regenerating clusters, and pink arrows indicate degenerated axons. Density of normal and degenerating axons **(I),** the average axon diameter **(J)**, and the distribution of axon diameters **(K)** in 12-month old WT and SCKO sural nerves. Density of normal and degenerating axons **(L)**, the average axon diameter **(M)**, and the distribution of axon diameters **(N)** in 12-month old WT and SCKO femoral nerves, n=3-4.

To further investigate the motor-selective neural deficits in LDH SCKO mice, we examined both sensory endings and motor synapses. We first assessed the integrity of sensory nerve endings in the skin by quantifying intraepidermal nerve fiber (IENF) density and found no difference in IENF density in 12-month-old mutant mice **(Fig. S7)**. In contrast, examination of the neuromuscular junctions in lumbrical muscles of the SCKO mice showed significant losses of innervated NMJs **(Fig. 7A-C)**. We also found that SCKO mice have smaller muscle fiber cross-sectional areas **(Fig. 7D-F)**, consistent with muscle atrophy secondary to partial denervation. In sum, these data strongly indicate that motor but not sensory neurons require LDH activity in SCs—and thus SC-derived lactate—to properly maintain their axons.

**Figure 7:**
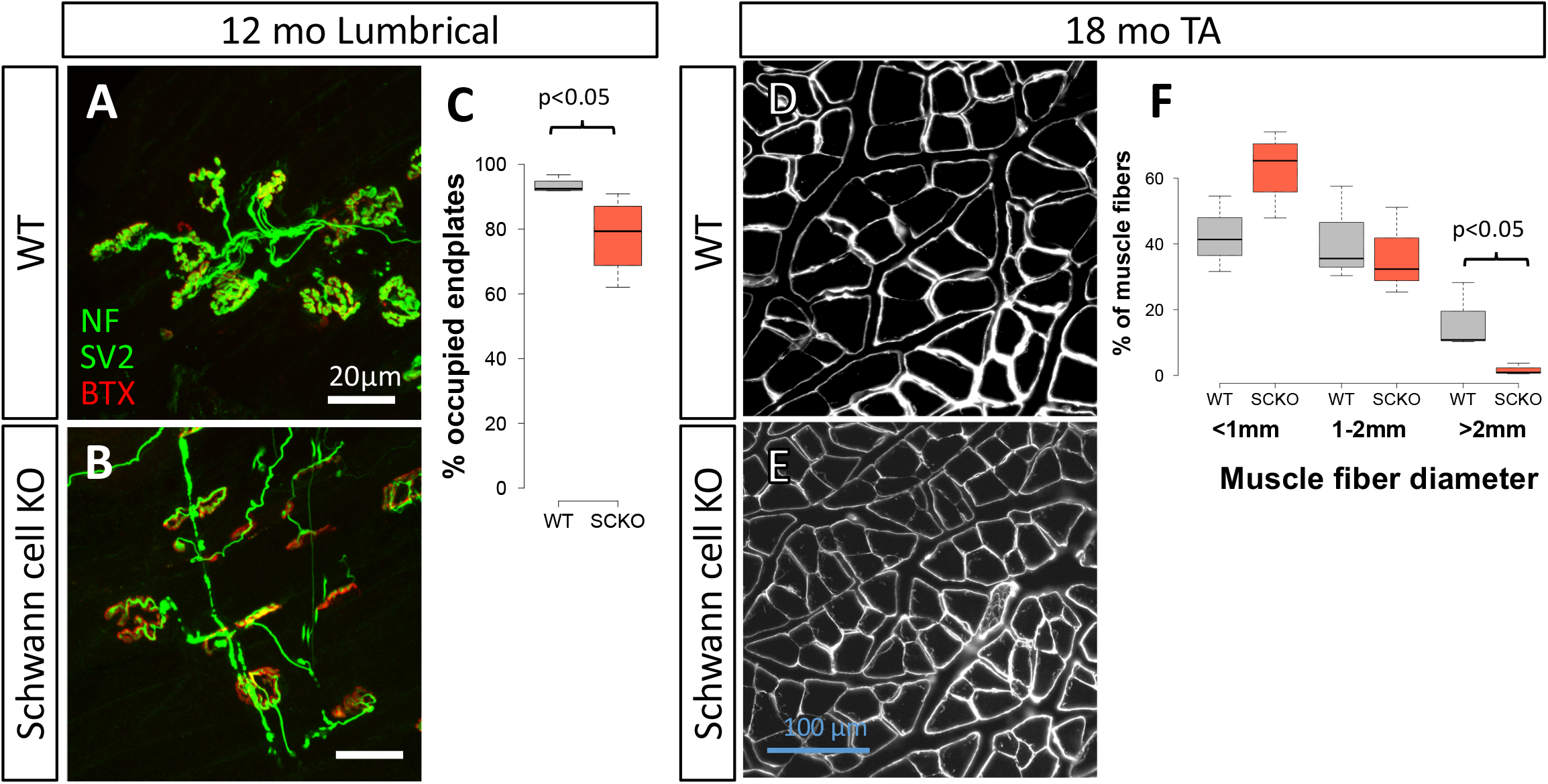
Muscle denervation and atrophy evident in SCKO mice. NMJs on 12-month-old control and SCKO lumbrical muscles stained in green to detect the synaptic vesicle marker synaptic vesicle glycoprotein 2A (SV2) and axon marker neurofilament medium chain (NF) and in red with the post-synaptic endplate ligand bungarotoxin (BTX) **(A,B)**. Percent of the lumbrical muscle endplates innervated by apposed pre-synaptic structures **(C)**. Representative images of tibialis anterior muscle cross-sections stained with laminin from 18-month-old WT and SCKO mice **(D,E)** with quantification of muscle fiber area **(F)**, n = 3-9 mice.

## Discussion

Levels of lactate production and lactate shuttling from glial cells to neurons in the CNS decline with age and are implicated in aging-related disorders including neurodegenerative diseases^39–41^. While growing evidence indicates parallel mechanisms in the PNS, the role of lactate metabolism in peripheral nerve degeneration remains undetermined. To explore the influence of Schwann cell (SC) lactate production on peripheral axon maintenance, we genetically targeted LDH activity (both the LDHA and LDHB isoforms) in either neuronal subtypes or SCs. Because LDH is responsible for interconverting pyruvate and lactate in both ‘lactate producer’ and ‘lactate consumer’ cells, LDH knockout in either cell effectively blocks lactate traffic between the axonal compartment and surrounding supporting cells. Surprisingly, we found that motor axons degenerate whether LDH activity is lost from either motor neurons or SCs, whereas sensory axons show little impact from loss of LDH either from sensory neurons or SCs. Overall, our results support a model in which motor axons are much more dependent than sensory axons on the lactate provided by SCs. This striking motor specificity implicates compromised lactate metabolism and shuttling as a potential pathogenic mechanism in motor-dominated neuropathies such as ALS.

Neurons require rapid and copious ATP production to support energetically demanding functions including propagation of action potentials, synaptic transmission, cytoskeletal maintenance, and axon transport ^42^. Cajal originally proposed that astrocytes directly supply nutrients to neurons^43^, and over the last quarter century evidence from multiple model systems has shown that glycolytic support cells specifically provision lactate to neurons to meet their extraordinary metabolic needs^9,44–49^ Once shuttled into axons, lactate can be converted into pyruvate and consumed by axonal mitochondria to produce ATP or building blocks for macromolecules. While lactate shuttling has been amply demonstrated in the CNS, PNS neurons also have high metabolic demands due to their large axons. Key observations indicate a parallel lactate shuttling pathway in peripheral nerves could play a critical role in peripheral neuropathy. For example, like astrocytes in the CNS, SCs in the PNS store glycogen whereas the axons they ensheath do not^11^. After nerve injury, SCs upregulate glycolytic metabolism to support axons and prevent their degeneration^50^. Mice deficient for monocarboxylate transporter 1 (MCT1)—which transports lactate, pyruvate and ketone bodies—in SCs develop spontaneous axonal defects^12,13^, indicating that SCs export a factor essential to axon health via MCT1. However, the MCT1 SC knockout phenotype is far milder than defects we observe in SC or motor neuron LDH knockout mice, suggesting additional mechanisms for lactate export from SCs exist. MCT4 is also expressed by SCs, but it appears largely dispensable for support of motor axons^48^. Connexin (CX32) may contribute to lactate transport from SCs to axons, as mutations in the CX32 gene associated with Charcot Marie Tooth disease type 1X reduce channel permeability to small molecules including lactate ^51–54^. CX32 mutant mice display motor-specific phenotypes remarkably similar to LDH SCKO mice^55,56^. Although CMT1X has been traditionally considered a demyelinating neuropathy, axonal changes including altered cytoskeleton and axonal transport precede demyelination in CMT1X, further highlighting the importance of the SC metabolic support of axons ^38^.

SC-specific disruption of metabolism leads to myelination defects and axon degeneration, principle symptoms of peripheral neuropathy^2–8,57^. In most of these mutant mice, sensory pathology predominates. For example, deletion of COX10 or TFAM, proteins important for mitochondrial metabolism, leads to hypomyelination and progressive axon loss^2–4^. Loss of INSR or IGF1R in SCs results in sensory-specific peripheral neuropathy^24^ and ablation of the central metabolic regulator LKB1 in SCs results in impaired myelination and progressive small-fiber axon degeneration^5,58^. LKB1-deficient SCs consume less oxygen and produce more lactate, a compensatory process that is axoprotective^5,58^. Conversely, when LDH is deleted in SCs, we see oxygen consumption increase and hypermyelination in the form of tomacula is detected at an early stage, with demyelination occurring subsequently. In addition to reduced lactate, nerves of LDH SCKO mice have reduced ATP, which may explain some of the axonal deficits and NMJ abnormalities^59^.

While motor and sensory axons apparently have different metabolic needs, SC subtypes that ensheath motor and sensory axons could also differ in their metabolism. Single nuclei and bulk RNAseq analyses of peripheral nerves from wildtype and SOD1^G93A^ ALS model mice identified a subpopulation of SCs that preferentially associate with motor axons and express elevated levels of LDH and other genes related to pyruvate and NAD metabolism. Crucially, this motor-associated SC population disappears as motor axon loss progresses in the SOD1^G93A^ model^60^.

A range of cellular processes are reported to influence liability for the most common motor neuron disease, ALS. These include aberrant RNA processing, ER stress, metabolic abnormalities, neuroinflammation, oxidative damage, and mitochondrial dysfunction, among others^61^. As with many neurodegenerative conditions, age is the primary risk factor for ALS, and there is clear overlap between pathways implicated in the disease and natural aging processes^62^. Age-related changes in metabolism—including lactate metabolism—are important feature of ALS^39,63^. For example, reduced levels of lactate and MCT1 are found in spinal cords of SOD1^G93A^ mice and motor cortex tissue of ALS patients carrying pathogenic SOD1 mutations, patient fibroblast-derived oligodendrocytes^64–66^, as well as in cultured astrocytes with TDP-43 inclusions^67^. Furthermore, lactate supplementation increases the survival of ALS model motor neurons *in vitro*^65^ and increases ATP production and survival in an in vitro model of glutamate induced neurotoxicity, a process associated with ALS pathology^68^. The expression of key rate-limiting enzymes in glycolysis (PFKP and PFKM) as well as flux through the pentose phosphate pathway (G6PD) are increased in spinal cord tissue and iPSC-derived neurons from ALS patients with TDP-43 proteinopathy^69^, findings that are similar to the accumulation of glycolytic intermediates and increased activity of G6PD in LDH SCKO animals **(Fig. 3, 3S).** While astrocyte and oligodendrocyte lactate shuttling are important for motor neuron survival^64^, SC lactate shuttling could be particularly important for maintaining distal motor axons and the NMJs. Indeed, in ALS models and patient autopsy tissue, NMJ pathology is correlated with the extent of motor deficits and axon degeneration precedes motor neuron cell death^70–72^. These findings highlight the importance of axon-SC metabolic coupling as a potential driver of peripheral ALS pathology.

Our unexpected discovery that disrupted LDH activity in SCs provokes motor-selective pathology reveals an essential difference between sensory and motor axons with implications for the pathogenesis and treatment of motor-dominated conditions including ALS. Only a few metabolic intervention strategies have been investigated in preclinical and early-stage clinical trials for ALS^63^. Therapeutic strategies to enhance glycolysis show some promise in models of ALS and other neurodegenerative diseases^73^, but given signs of glycolytic compensation in human neurons with ALS pathology^69^, and the dependence of motor neuron health on lactate metabolism, it is reasonable to consider targeting cellular metabolism in a manner that restores mitochondrial ATP production while circumventing impaired glycolysis^74^. Treatments that seek to replace glucose as a cellular fuel source—such as through supplementation with pyruvate, lactate and other substrates—or to otherwise ramp up TCA cycling and mitochondrial oxidative phosphorylation, have shown some success (reviewed in ^63,74–76^). Ultimately, metabolic strategies that account for the lactate-dependence of motor axons and NMJs may be useful in targeting dysregulated metabolism in ALS.

## Acknowledgments

We would like to thank members of the DiAntonio and Milbrandt labs for their thoughtful feedback on this work. We would like to thank Cassidy Menendez, Rachel McClarney, Liya Yuan, and Alicia Neiner for their technical support. We also thank the Washington University Core for Cellular Imaging (WUCCI) for their technical support, expertise, and training. We also thank the Genome Engineering & Stem Cell Center (GESC@MGI) at Washington University for generating transgenic mice.

---

**Supplemental Figure S1:** Measurement of Compound Muscle Action Potential (CMAP) at the Sciatic Notch Amplitude **(A,B)** and latency **(C)** in MNKO mice. Sensory Nerve Action Potential (SNAP) amplitude and conduction velocity (CV) measured from the tail **(H,I)** in SNKO mice.

**Supplemental Figure S2:** To assess metabolic flux, 3-month-old sciatic nerves were incubated in glucose-free media supplemented with ^13^C_6_ glucose for 24 hours *ex vivo*. Total lactate detected **(A)**. Diagram showing carbon isotope labeling of glucose and its production of labeled pyruvate and lactate **(B)**. (Total lactate measured in M+0 (Unlabeled), M+1, M+2, M+3 (3 carbons labeled, the direct product of glycolytic pyruvate) **(C)**. *p<0.05 using a Student’s t-test between genotypes for (B-C). *p<0.05 using a Student’s t-test between genotypes for each gene (A). n = 3-12 mice.

**Supplemental Figure S3:** Metabolic analysis the pentose phosphate pathway **(A-D)**, and purine metabolism **(E-H)**. Metabolites in gray were undetected, metabolites in black were unchanged, and metabolites in red were upregulated. n=3-5 mice. *p<0.05 using a Student’s t-test between genotypes at each time point. **(I)** The activity of Glucose-6-phosphate dehydrogenase (G6PDH), the rate-limiting step in the pentose phosphate pathway, is significantly increased in SCKO mice. n=3-5 mice. *p<0.05 using a Student’s t-test between genotypes at each time point.

**Supplemental Figure S4: (A-D)** Toluidine blue-stained sciatic nerves from WT and SCKO mice at postnatal day 7 and 21. Quantification of the number of myelinated axons **(E)** or Schwann cell nuclei **(F)** in WT or SCKO mice at postnatal day 7 and 21. n=3 mice. Scale = 20 μm.

**Supplemental Figure S5: (A)** Toluidine blue-stained sciatic nerves from WT and SCKO mice at 14 days after sciatic nerve crush injury. Quantification of the number of myelinated axons **(A)**, naked axons **(B)**, nuclei **(C)**, or foamy macrophages **(D)** in WT or SCKO mice at postnatal day 7 and 21. n=3-4 mice. Scale = 20 μm.

**Supplemental Figure S6: (A-F)** Whole nerve cross-sections of toluidine blue-stained proximal sciatic nerves from WT and SCKO mice at 3, 6, and 12 months of age. Scale = 100 μm. **(G-L)** Zoomed in images of proximal sciatic nerves from WT and SCKO mice at 3, 6, and 12 months of age. Orange arrows indicate tomacula. Yellow arrows indicate naked axons. Pink arrows indicate degenerated axons. The Blue arrow indicates an onion bulb. The green arrow indicates a regenerating cluster. Scale = 20 μm.

**Supplemental Figure S7:** Sensory nerve endings do not degenerate in SCKO mice. **(A-D)** Representative images of WT and SCKO mouse footpad skin stained with the pan-axonal marker PGP9.5 (red) and the nuclear marker DAPI (blue). The dashed white line shows the basement membrane. Scale = 200 μm. **(E)** Quantification of intraepidermal nerve fiber (INEF) density per mm of basement membrane. n = 3-9 mice. *p<0.05 using a Student’s t-test between genotypes for each category of innervation.

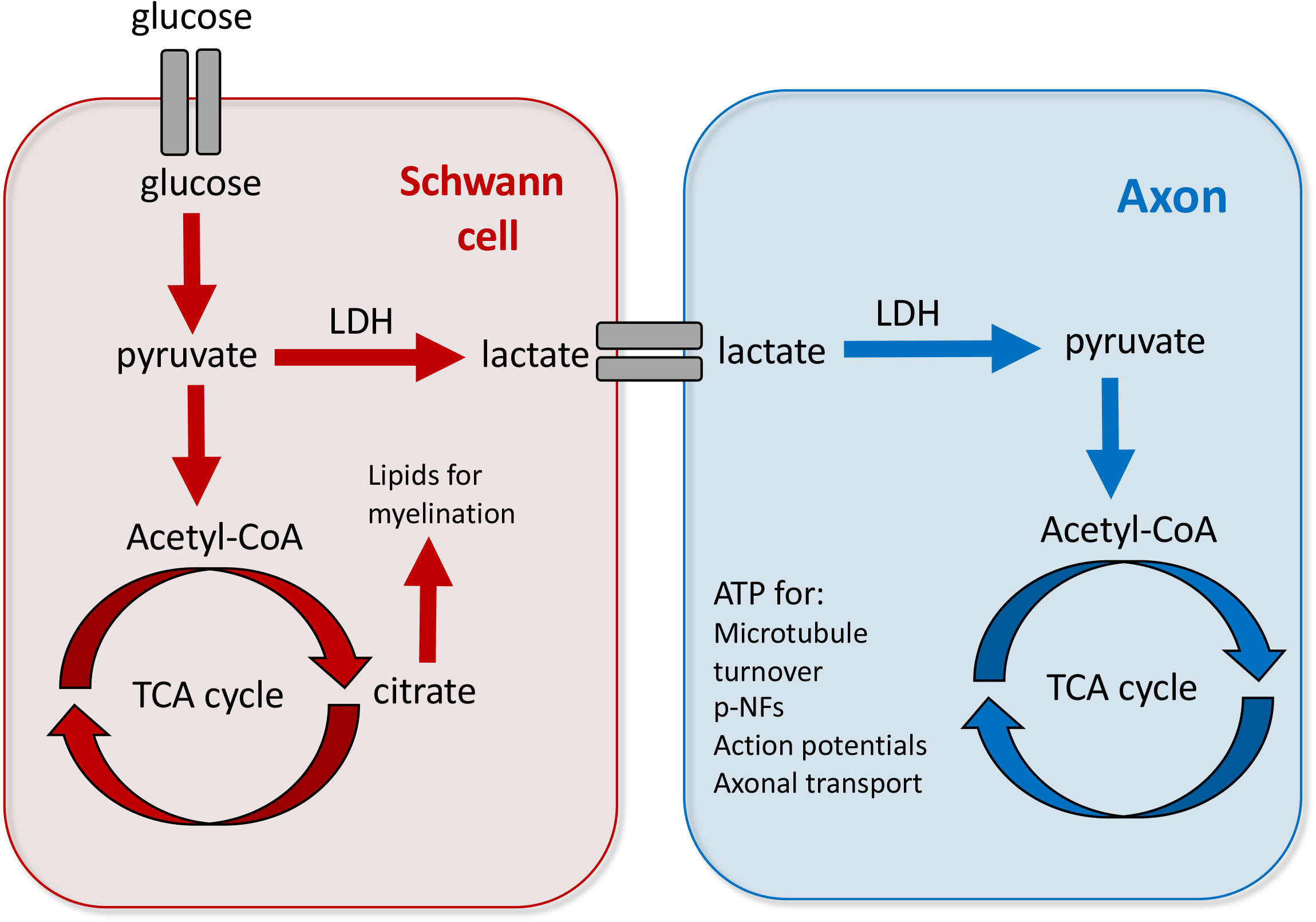
Graphical Abstract

